# Kinship, acoustic signaling, and socio-spatial structure shape shared decisions and social influence in white-nosed coatis

**DOI:** 10.64898/2026.08.11.744150

**Authors:** Jack C. Winans, Emily M. Grout, Josué Ortega, Matthew J. Quin, Margaret C. Crofoot, Ben T. Hirsch, Ariana Strandburg-Peshkin

## Abstract

When individual preferences for collective outcomes diverge, cohesive animal groups often coalesce on the majority opinion. However, majority-based decision rules may be counterbalanced by other factors, particularly in heterogeneous groups with differentiated social relationships, and these factors could produce inequality in social influence. We used multi-sensor tracking collars to collect detailed and simultaneous data on the movements and vocalizations of almost all members of three wild white-nosed coati (*Nasua narica*) groups, and analyzed 2,401 individual decisions between conflicting travel directions that were indicated by acoustically signaling groupmates. Decision-making was shared: individuals favored directions that had majority support, and we found evidence that they used acoustic signals and movement cues to infer majority support. Individuals were also more likely to choose directions favored by closer kin and by groupmates in more frontward spatial positions. Although decisions were shared, influence was not equally distributed across individuals. Individuals who consistently formed majorities or occupied frontward spatial positions had higher influence over travel direction. By explicitly linking decisions by individual followers to emergent patterns of influence among potential leaders, our results suggest that influence is a complex product of higher-order interactions that are likely dependent on group demography and socio-spatial structure.

## Introduction

Animal groups often include members with different energetic requirements [1], locomotor capacities [2], personal information about the environment [3], competitive abilities [4], and susceptibility to predation [5]. These differences can lead to conflicts of interest over when to exploit which resources [6]. If groupmates fail to overcome their conflicting interests and reach consensus about travel direction and timing, they risk losing the benefits of group cohesion [7, 8]. A growing body of evidence suggests that animal groups commonly reach consensus via shared decision-making processes that integrate the preferences of many or all group members (e.g., preference averaging [9, 10], quorum-based decisions [11–13], or majority-based decisions [7, 9, 10, 14]; see table 1). However, it is unclear how complex social dynamics and differentiated relationships impact these processes in heterogeneous groups [15]. As such, understanding how social interactions shape the processes by which groups reach consensus, and ultimately shape the consequences of consensus decisions, remains a central goal of the study of animal collective behavior.

**Table 1.** Definitions of collective behavior-related terms.

| Term | Definition | Reference |
|---|---|---|
| Consensus decision | A collective process in which all members of a group converge on one of multiple, mutually exclusive behavioral outcomes, either through a shared, partially-shared, or unshared process | [8] |
| Shared decision | A consensus decision that integrates the outcomes preferred by many or all group members. | [8, 56] |
| Unshared decision | A consensus decision that only takes the preferred outcome of a single individual into account. | [8, 56] |
| Social influence | In general, social influence is the propensity for the actions of one individual to result in a behavioral change among other individuals. In this study, social influence is quantified by the probability of attracting groupmates to one’s own preferred travel direction, with this preference inferred based on spatial position and contact calling behavior. | [56] |
| Higher-order interaction | A set of simultaneous social interactions among three or more individuals that cannot be reduced to or explained by dyadic interactions among the contained individuals. | [17, 54] |

Consensus decisions reached via shared processes are frequently contrasted with those reached via unshared processes [8, 16]. Unlike shared decisions, when conflicts arise that could jeopardize group cohesion, unshared decisions entail all group members conforming to a leader’s preference [8]. Nevertheless, whether consensus decisions are shared or unshared, they are similarly built up from many individual decisions by group members to either pursue their own preferred outcome or forego it for another’s.

Although these decisions (to pursue or forego one’s preference), and their individually-specific and socially-mediated drivers, are the source of variation in social influence in heterogeneous groups [10, 17, 18], studies on animal collective behavior rarely link the multiple factors weighed in decisions by potential ‘followers’ to the emergence of particular ‘leaders’.

Explicitly testing this relationship between the individual decision-making process and group-level patterns of social influence has the potential to clarify the mechanisms and consequences of consensus decisions. In particular, understanding the links between the decision biases of individuals and the choice made by their group as a whole could reveal how interindividual differences in social influence can emerge, even when consensus decisions are mechanistically shared.

Group movements are conspicuous collective decisions in which individuals can ‘vote with their feet’ by moving toward a preferred travel direction [19, 20]. Theoretical models and empirical data on animal movement trajectories support a ‘compromise or choose’ pattern during shared directional decisions: when the directions preferred by different groupmates are relatively similar, groups tend to ‘compromise’ by averaging the preferred directions, but when directions preferred by different groupmates are relatively dissimilar, groups instead ‘choose’ the direction preferred by the majority of individuals [7, 9, 10, 21–23]. Instances of high directional disagreement therefore offer unique insight into the individual decision-making processes that structure social influence, because they offer obvious examples of individual choice (from the perspective of potential followers) and of being chosen or not (from the perspective of potential leaders).

Individual trajectories can be used to detect ‘votes by feet’ in a shared directional decision, but, in reality, such decisions are often coordinated by complex exchanges of multimodal signals and cues [24, 25]. Because individuals likely base their decisions on social information perceived through multiple senses, even high-resolution positional tracking can be limited in its ability to reconstruct decision-making processes during directional disagreement. Multi-sensor tracking tags can provide synchronized data streams on both individual movements and vocal signals [26–28], allowing for more detailed reconstructions of the sensory landscape shaping decisions.

Here, we used multi-sensor tracking tags to study collective decisions about travel direction in heterogeneous social groups in the wild. We investigated the process of directional conflict resolution at two scales: individual directional decisions by followers and aggregate levels of social influence by potential leaders, and attempted to link these scales. At the level of individual follower decisions, we identified multimodal factors predicting directional choice. At the aggregate level, we tested whether individuals exhibited consistent differences in social influence over directional decisions. Finally, we tested whether the predicting factors identified at the level of individual follower decisions could explain variation in social influence at the aggregate level.

We tracked the movements and vocalizations of all, or almost all, members of three wild groups of white-nosed coatis (*Nasua narica*), diurnal procyonids ranging from northern Colombia to the southwestern United States across diverse forested habitats [29, 30]. Adult females and their dependent offspring live in social groups with relatively stable membership (four to ∼30 members) that sometimes split into smaller subgroups as they forage for leaf-litter invertebrates and ripe fruit [30–33]. When groups split, kinship often explains individual subgroup choice, such that relatedness tends to be higher within than between subgroups [34]. Males disperse from their natal group shortly before sexual maturity and typically live alone as adults, but associate with female-bonded groups during the brief mating season [30, 33, 35]. Coatis also possess a varied vocal repertoire, and emit short-duration, high-frequency contact calls while on the move [36–38].

We tested three main hypotheses. The first two of these hypotheses were specific to the level of individual follower decisions, and the third was related the emergence of social influence among potential leaders. First, because coatis often live in densely vegetated habitats and likely evolved from a recent nocturnal ancestor with poor vision [39, 40], we hypothesized that acoustic signaling facilitates shared directional decisions by coati groups (H1). To test H1, we evaluated three potential proxies by which coatis could perceive a majority opinion via movement cues, acoustic signals, or both. We used these three proxies to test our prediction that individuals would be more likely to choose the travel direction indicated by a majority of their groups (P1.1). We then compared the extent to which each of these proxies predicted directional choice to test our prediction that vocal majorities (i.e., greater spatial concentrations of contact calls) would be more predictive of individual directional decisions than absolute majorities based on all group member locations (P1.2).

Second, we hypothesized that individuals ‘weigh’ the preferences of close social affiliates as much as, or possibly more than, they ‘weigh’ the majority opinion (H2). We predicted that individuals would be more likely to choose the travel direction indicated by a more closely-related groupmate (compared to groupmates indicating the alternative direction; P2.1), given their strong tendency to associate with more closely related groupmates [34]. Next, we compared the strength of the tendency to follow the majority opinion (tested under P1.1-2) versus the strength of the tendency to follow closer kin, and predicted that they would have comparable effects on individual directional choice (P2.2).

Our third hypothesis was that the mechanisms of decision making at the individual follower level (tested under H1-2) would produce individually-consistent variation in social influence over group travel direction (H3). To test H3, we predicted that when these conflicts occurred, groups would be more likely to move toward directions indicated by particular individuals (i.e., those with high social influence) than toward directions indicated by some other individuals (i.e., those with low social influence; P3.1). Finally, we predicted that these differences in social influence would be explained by the factors predicting individual directional choice investigated under H1 (i.e., individual tendencies to form majorities and to be more closely-related to potential followers; P3.2).

## Methods

### Study site and subjects

We sampled three social groups of white-nosed coatis in two semideciduous lowland tropical forest sites in Panama: Barro Colorado Island (9°16′N, - 79°83′W) and Soberania National Park (9°12′N, -79°70′W). These sites are approximately 16 km apart and have been separated by the waters of Lake Gatun since 1914, following the damming of the Chagres River. The region is characterized by a dry season that runs from late December or early January until the start of the wet season in late April or early May [41].

Two of our study groups, Galaxy group and Trago group, were sampled in the dry season of 2021-2022 in SNP, and the other, Presidente group, was sampled in the dry season of 2023 on BCI. Because births are highly synchronous within groups and occur annually, we could assign individuals to one of three age classes with high certainty based on body size and secondary sex characteristics, following Gompper [31]: ‘juveniles’ (< 12 months old), ‘sub-adults’ (12-24 months old), and ‘adults’ (> 24 months old). In this population, the mating season occurs during the dry season, but its exact timing varies between social groups [34]. Each group was sampled at a different time relative to its mating season, which led to some of the between-group differences in demography. Presidente group was sampled prior to the start of the mating season and included three adult females, seven sub-adults, and six juveniles (16 total individuals).

Galaxy group was sampled during the mating season, meaning that, in addition to eight adult females, three sub-adults, and one juvenile, an adult male associated with the group (12 total individuals). Trago group was sampled after the mating season, when all adult females had left the group to give birth alone [33], and this group included one injured sub-adult male and six juveniles (seven total individuals).

### Data collection

We captured individuals with Tomahawk traps and immobilized them with Telazol (50 mg/mL tiletamine and 50 mg/mL zolazepam; 5.40 ± 0.50 mg/kg). We fitted each captured individual with a custom-built collar that had an onboard Global Positioning System (GPS) sensor (e-obs GmbH, Grünwald, Germany) and two miniature audio recorders (Soroka 18E, TS-Market Ltd., Zelenograd, Russia). All members of Presidente and Trago groups were collared, and 11/12 members of Galaxy group were collared (one adult female was not collared). See Supplementary Methods and Grout *et al*. [34] for additional collaring details.

The three tracking periods lasted from 24 December 2021 to 13 January 2022 (Galaxy), 24 March 2022 to 10 April 2022 (Trago), and 19 January 2023 to 2 February 2023 (Presidente). Collars were programmed to record one GPS location every second from 0600 to 0900. Mean GPS fix success rates were 96.5% (Galaxy), 96.1% (Trago), and 96.9% (Presidente) and mean GPS error (inferred by positioning collars at known distances from one another in the study area prior to deployment) was 3.86 m ± 1.06 s.d. The collar-mounted audio recorders collected data from 0600 to 0900 at a 24,000 Hz sampling frequency with 0 dB gain and 16-bit resolution.

Because we expected that the battery life of the audio recorders would deplete faster than the battery life of the GPS sensors, we programmed one audio recorder on each collar to start collecting data on the first day of the tracking period and the second audio recorder to start collecting data seven or eight days later. On average, audio recorder batteries lasted 8.12 days ± 0.70 s.d. We also anticipated that the internal clocks contained in each audio recorder would drift slightly over the course of the deployments. To ensure that GPS data and audio data could be accurately time-matched, we systematically played synchronization sounds in the field at known times during data collection periods, in the detection range of the deployed audio recorders.

### Ethics

We followed the ethical guidelines of the American Society of Mammalogists and the Smithsonian Institutional Animal Care and Use Committee for all animal capture and collaring procedures (IACUC clearance number: 2017-0815-2020). In all cases, collars weighed < 5% of subject body weight. See Grout *et al*. [34] for detailed information on efforts taken to minimize study subject stress and injury risk.

### Data processing

Full details regarding GPS and audio data processing performed prior to formal analyses are given in the Supplementary Methods. In brief, we trained a self-supervised transformer-based neural network model (*animal2vec*, [42]) to detect and classify coati vocalizations from a subset of labelled audio data. For the purposes of this study, we were interested in call types hypothesized to serve as ‘contact calls’: *chirp*, *chirp click*, *chirp grunt*, *click*, and *click grunt* [38, 43]. These calls are thought to coordinate movement because they are usually produced while coatis are moving and are not obviously associated with other social functions, such as aggression [38, 43]. Based on an initial training/validation split of the labelled data set and subsequent manual verification of detected calls, we estimated that focal contact calls (i.e., those that were emitted by the animal wearing the collar) were detected with a precision of 0.81 and a recall of 0.81 (figure S1; see *GPS and audio data processing* in Supplementary Methods). To time-match each detected contact call to the GPS data, we listened to the audio files to identify the time in each file that a synchronization sound was audible. We used the known times of these synchronization sounds to align the audio file time with GPS time and calculate contact call rates for each individual across the study period (see Supplementary Methods for how rates were calculated).

### Defining influence contests

We sought to explain how conflicts of interest over travel direction were resolved and how this conflict resolution process produced individual differences in social influence. Therefore, we combined our spatial and audio data to identify events in which a stationary individual chose to move in one of two mutually exclusive directions that were each simultaneously indicated by the presence of groupmates giving contact calls. As shorthand, we hereafter refer to these events as ‘*influence contests*’ and to the stationary individual who chose one direction as the contest’s ‘*decider*’. Because each directional option could have been indicated by several individuals, we refer to each collection of individuals indicating a single directional preference as a ‘*set*’ of ‘*contest participants*’ (figure 1 illustrates an influence contest example from our data set).

**Figure 1.**
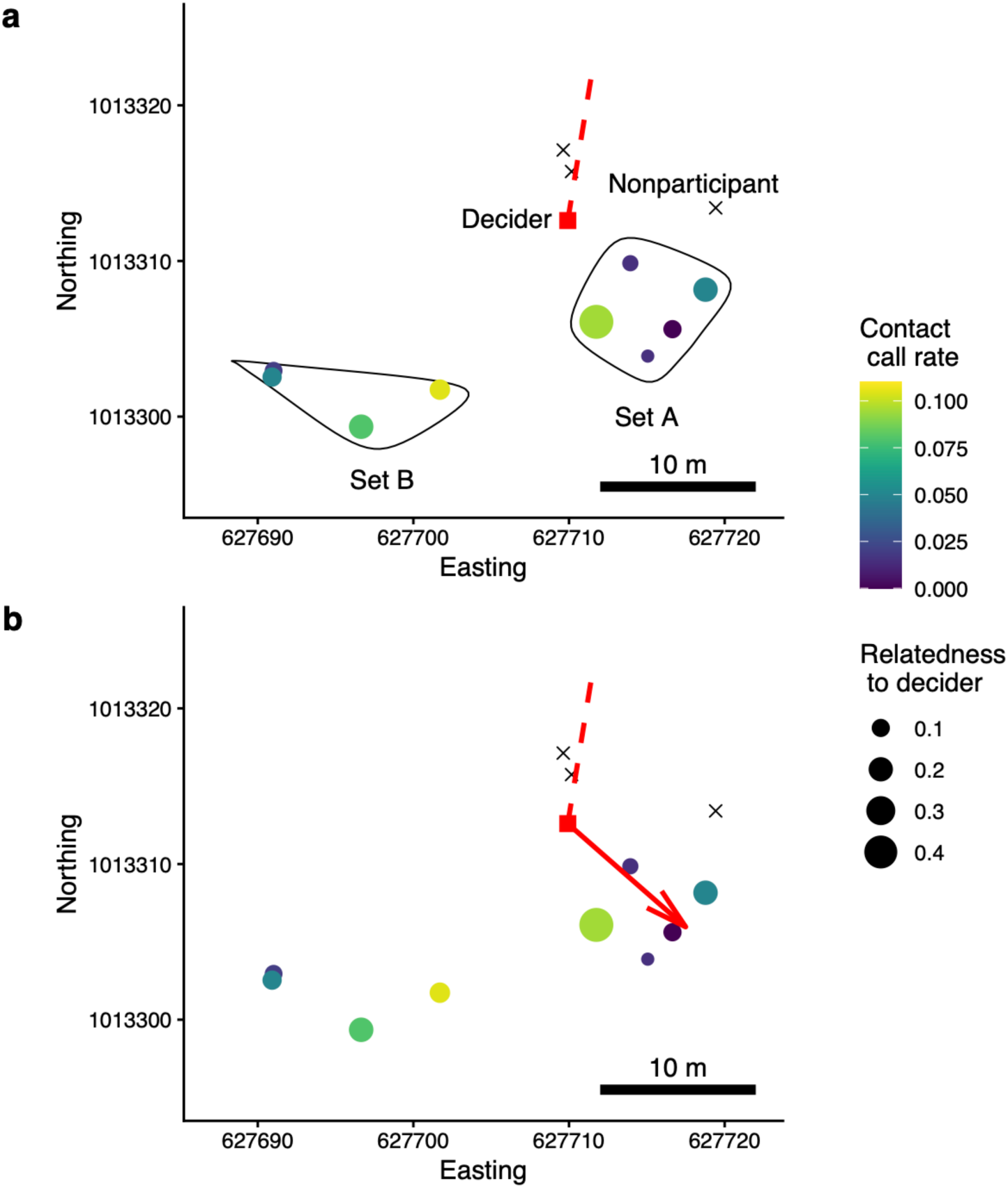
Example of an influence contest. These figures depict a randomly selected influence contest in Presidente group (three Presidente members were in a different subgroup at this time and are not shown here). In both panels, every point (circles, squares, and crosses) depicts the spatial position of one coati at time *t*. The red square represents the contest’s decider, the circles depict set participants, and the crosses represent nonparticipants (see Supplementary Methods for definitions). The sizes of participants’ circles represent their relatedness to the decider. Circles are colored according to the participants’ contact call rates at time *t*. The dotted red line shows the past heading of the decider (moving southwestwardly). In (a), the two identified sets (set A and B) are enclosed with a black line. In (b), the solid red line shows the decider’s future heading (moving southeastwardly in the direction proposed by set A).

The utility of this approach is twofold. First, from the perspective of contest deciders, we can estimate the relative strengths of different biases involved in making complex social decisions (H1-2). Second, from the perspective of contest participants, we can measure interindividual differences in the probability of ‘*winning*’ (i.e., attracting the decider), which should approximate differences in social influence over travel direction (H3).

In our analyses, each influence contest was defined as a discrete event during which two sets of coatis were giving contact calls from different directions (separated by an angle > 90°) relative to a stationary decider, who then moved at least 10 m in one of those directions at > 0.2 m/s. By limiting our definition of influence contests to cases where at least some members of both sets were producing contact calls, we intended to restrict our data set to cases in which participants in both sets had some movement-related motivation (coatis tend to emit contact calls while they are moving and increase their contact call rate with increasing speed [43]). A complete description of the analytical method used to detect these events and of our validation of these detected contests can be found in the Supplementary Methods.

### Set-level advantages during influence contests

We calculated proxies of a set’s relative advantage over its opposing set in three domains: numerical support, kinship, and frontness (figure 1). A complete description of the rationale underlying these sources of advantage and of the methods used to calculate them can be found the Supplementary Methods. The advantage of a given set in a particular domain could range from -1.00 to 1.00, with negative values indicating that the set was relatively disadvantaged and positive values indicating that the set was relatively advantaged in a given domain.

We calculated three separate proxies of numerical support advantage to account for potential variation in the extent to which coatis rely on acoustic signals versus cues about movement (visual, olfactory, or auditory indicators of neighbor locations and travel directions) to infer which direction most of their groupmates are going [25]. These proxies included (i) *presence advantage* (positive values indicate that the set had more total members than its opposing set), (ii) *signaler count advantage* (positive values indicate that the set had more members producing contact calls than its opposing set), and (iii) *signal rate advantage* (positive values indicate that the set, as a whole, was producing contact calls at a higher rate than its opposing set). We also calculated a set’s (iv) *kinship advantage* (positive values indicate that the decider’s closest relative on the set was more closely related to the decider than its closest relative on the opposing set) and (v) *frontness advantage* (positive values indicate that the set was relatively farther in front of the decider than its opposing set).

*H1-2: Factors shaping individual movement decisions during instances of high directional disagreement.* To test our first two hypotheses, we randomly selected one set in each detected influence contest to serve as the focal set. We constructed three Bradley-Terry models for each group [44, 45], as Bernoulli generalized linear mixed models (GLMMs) with logit-link functions and signed multi-membership random effects. Commonly used to predict the probability that an individual member of a team wins a paired match, given that team composition fluctuates between matches [46], Bradley-Terry models include a random effects matrix with a positive indicator (+1) for each member of the focal set, a negative indicator (-1) for each member of the opposing set, and a neutral indicator for any individuals who do not participate in a given contest. Thus, a random intercept can be estimated separately for each individual, based on all contests in which they participated in either the focal or nonfocal set.

In each model, a trial consisted of a single influence contest and the response variable was whether or not the randomly-selected focal set won the influence contest (i.e., attracted the decider). In addition to the random effects matrix accounting for baseline interindividual variation in the probability of winning, each model included three fixed effects: (i) the focal set’s kinship advantage, (ii) the focal set’s frontness advantage, and (iii) either the focal set’s presence advantage, signaler count advantage, or signal rate advantage. In the model that included presence advantage, the random effects matrix included all members of both sets. In the signaler count and signal rate advantage models, the random effects matrix included all members of both sets whose contact call rate at the time of the influence contest was > 0.00 calls/s. To compare the three alternate model structures within each group, we calculated Akaike’s information criterion (AIC) for each model. When the ΔAIC between two models was > 2.00, we considered that the model with the lower AIC value was a better fit to the data.

*H3: Individual variation in social influence.* For each individual, we identified every influence contest in which they participated as a signaling member of a set. Multiple influence contests sometimes clustered in time when several deciders were choosing between the same, or very similar, sets. This becomes problematic when taking the perspective of contest participants, rather than deciders, because successive contest outcomes are likely non-independent. Therefore, we split our data set of influence contests into 5-minute intervals. For each individual, we randomly sampled a single influence contest in which they participated as a signaling member of a set from each 5-minute interval (if they participated in any during that interval). Contest outcomes in the resulting data sets were not significantly temporally autocorrelated, indicating this method resolved the statistical non-independence issue.

From these individual-based, temporally subsampled data sets of influence contests, we calculated a *realized* influence score for each individual as the observed proportion of influence contests that they won. For each individual, we also generated a null distribution of influence scores that would have been expected if deciders chose sets randomly with respect to the identity of the participant of interest. We considered that an individual had higher than expected social influence if their observed influence score was above the 97.5^th^ percentile value of their null distribution and that an individual had lower than expected social influence if their observed influence score was below the 2.5^th^ percentile value of their null distribution.

To assess whether social influence was related to the biases of deciders evaluated for H1-2 (i.e., biases toward majorities, kin, and frontal spatial positions), we took two complementary approaches. First, we took advantage of the random effects structure of the GLMMs used to test H1-2 to estimate a *baseline* influence score for each individual. This baseline influence score was calculated as the model-predicted probability that an individual would win an influence contest, holding each of the predicted advantages (presence/signaler count/signal rate, kinship, and frontness) constant at zero. If interindividual differences in realized social influence chiefly resulted from the biases of deciders, then these baseline influence scores should show little interindividual variation and converge around 0.50. The decision to use the GLMM including presence, signaler count, or signal rate advantage to calculate baseline influence scores was made separately for each group and based on the results of our test of H1.

Next, we calculated the mean value of each advantage type experienced by each individual across the subsampled influence contests in which they were a signaling set participant. For each group, we constructed a binomial generalized linear model (GLM) with a logit-link function predicting the proportion of influence contests that each individual won. We included three fixed effects in this model: (i) the individual’s mean kinship advantage, (ii) the individual’s mean frontness advantage, and (iii) either the individual’s mean presence advantage, mean signaler count advantage, or mean signal rate advantage. Again, the decisions to use mean presence, signaler count, or signal rate advantage in this model was made separately for each group and based on the results of our test of H1.

### Software

All analyses were conducted in R v4.5.0 [47] and statistical models were constructed with the *lme4* and *lmerMultiMember* packages [48, 49]. R code and data necessary to replicate all statistical analyses can be found in [50].

## Results

### Influence contests

We detected 2,401 unique decisions by individual coatis between mutually exclusive travel directions indicated by signaling groupmates (n = 1,596 decisions by 16 individuals in Presidente group, n = 732 decisions by 11 individuals in Galaxy group, n = 73 decisions by 7 individuals in Trago group). The relative scarcity of influence contests in Trago group was not surprising, given that their data were collected in the absence of all adult females (who had left the group to give birth just before data collection). Trago group was consistently spatially cohesive and rarely moved substantial distances during the study period, possibly because juveniles may be at higher risk of predation than larger adults. We present all results for Trago group in the Supplementary Material. These results were generally similar to the results for Presidente and Galaxy groups, but we caution that the small sample size limits further interpretation. Further summary statistics regarding detected influence contests, and their consequences for group-level movement, can be found in the Supplementary Results.

### H1-2: Movement decisions during instances of high directional disagreement were predicted by majority opinion, kinship, and spatial positioning

The members of Presidente group and the members of Galaxy group seemed to differ in the ways in which they integrated movement cues and acoustic signals to inform their directional decisions. In Presidente group, the probability that a focal set attracted a decider was better predicted by the set’s presence advantage than by its signaler count advantage (ΔAIC = 21.60) or signal rate advantage (ΔAIC = 27.52). Galaxy group exhibited a different pattern: the model including signal rate advantage had the lowest AIC value, but the focal set’s signal rate advantage was similarly predictive of the its probability of attracting a decider to its presence advantage (ΔAIC = 1.84) and its signaler count advantage (ΔAIC = 1.48). For each of these two groups, we present the results of the model with the lowest AIC value in the main text, and the results of the other models in tables S1-4. In each model, the coefficient estimate for the fixed effect approximating perceived majority opinion (presence advantage, signaler count advantage, or signal rate advantage) was positive and statistically significant, suggesting general support for majority-based decision making in coatis (P1.1), regardless of the mechanisms by which individuals might perceive conspecific preferences.

In Presidente group, a given set was more likely to attract a decider when it contained a relatively higher number of total individuals compared to its opposing set (P1.1-2; β = 1.11, *p* = 2.2 x 10^-8^; table 2, figure 2a). Directional choice was also predicted by the difference in relatedness between the decider and its closest relative in each set: deciders were more likely to choose the set containing the closer relative (P2.1; β = 0.72, *p* = 5.7 x 10^-5^; table 2, figure 2b). A set that was farther in front of the decider than its opposing set was also more likely to attract the decider (β = 0.95, *p* = 6.6 x 10^-7^; table 2, figure 2c).

**Figure 2.**
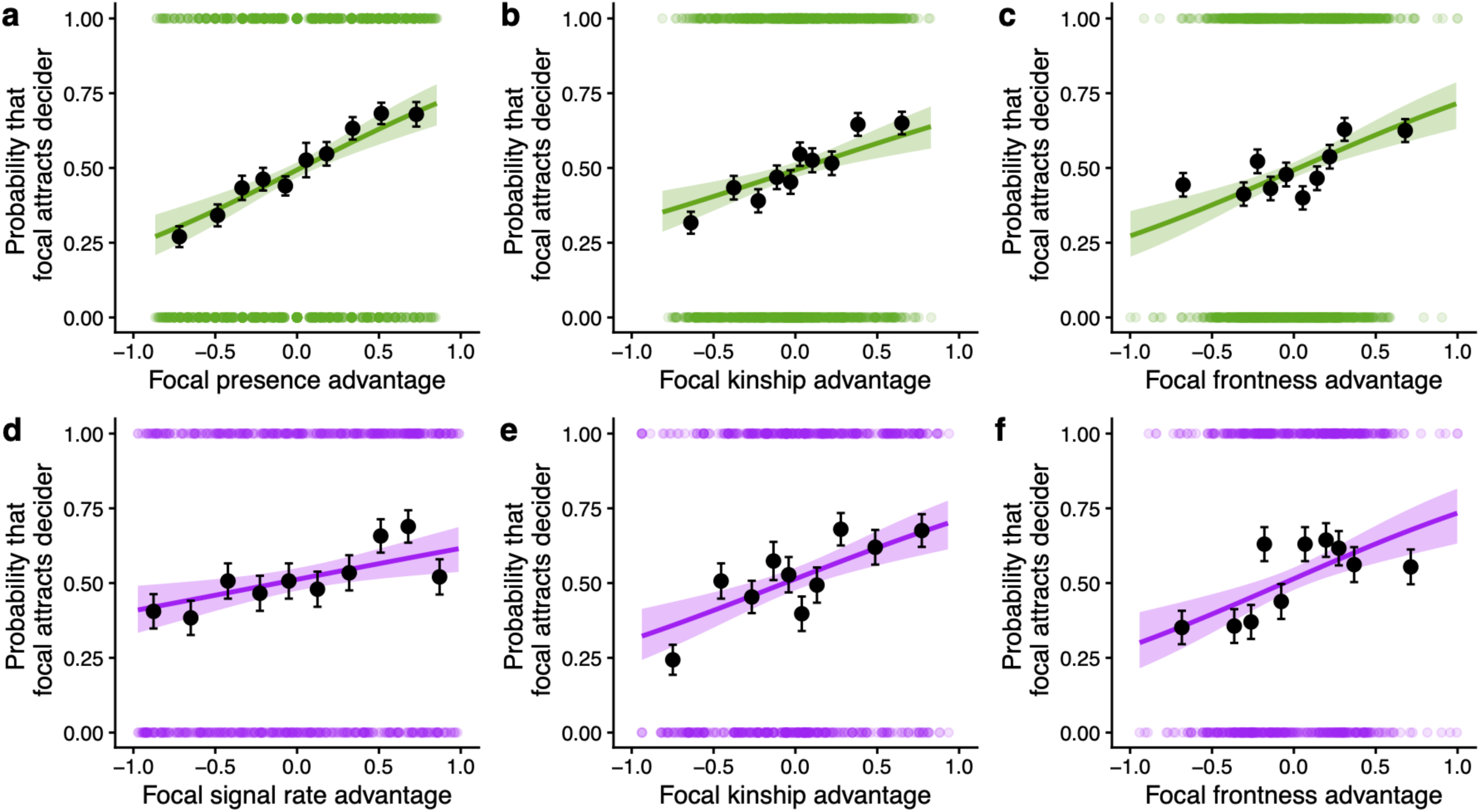
Majority opinions, kinship, and spatial positioning shape directional decisions during influence contests. Focal sets were more likely to attract deciders when they had a numeric or vocal majority (a,d), contained a signaling member who was more closely related to the decider than the decider’s closest relative on the opposing set (b,e), and when they were in a more forward position, relative to the decider’s past heading (c,f). (a-c) depict results for Presidente group and (d-f) depict results for Galaxy group. In each panel, each colored dot (green or purple) represents a single contest outcome (0 = focal set lost, 1 = focal set won). The black dots each represent the mean probability of winning observed for one decile of data, and the error bars depict the standard errors of these means. The colored lines and surrounding shaded regions depict the model-predicted relationships and their 95% confidence intervals.

**Table 2.** Results of Bernoulli GLMMs predicting the probability that a randomly selected set of participants in an influence contest successfully attracted a decider in Presidente group (n = 1,596 decisions by 16 individuals) and Galaxy group (n = 732 decisions by 11 individuals). Statistically significant fixed effects are indicated in bold. Here, we present the results for the Presidente model that included focal presence advantage and results for the Galaxy model that included focal signal rate advantage (see tables S1-4 for results of models with alternative proxies for majority opinion).

| Group | Fixed effect | Estimate | Std. Error | *p*-value | Interpretation |
|---|---|---|---|---|---|
| Presidente | (intercept) | -0.03 | 0.05 | 0.59 | – |
|  | **Focal presence advantage** | **1.11** | **0.20** | **2.2 x 10<sup>-8</sup>** | 1 s.d. increase in advantage = 24.48% increase in probability of attracting decider |
|  | **Focal kinship advantage** | **0.72** | **0.18** | **5.7 x 10<sup>-5</sup>** | 1 s.d. increase in advantage = 12.20% increase in probability of attracting decider |
|  | **Focal frontness advantage** | **0.95** | **0.18** | **6.6 x 10<sup>-7</sup>** | 1 s.d. increase in advantage = 14.17% increase in probability of attracting decider |
| Galaxy | (intercept) | 0.04 | 0.08 | 0.62 | – |
|  | **Focal signal rate advantage** | **0.42** | **0.15** | **4.5 x 10<sup>-3</sup>** | 1 s.d. increase in advantage = 11.26% increase in probability of attracting decider |
|  | **Focal kinship advantage** | **0.85** | **0.23** | **3.4 x 10<sup>-5</sup>** | 1 s.d. increase in advantage = 16.97% increase in probability of attracting decider |
|  | **Focal frontness advantage** | **0.85** | **0.20** | **2.0 x 10<sup>-5</sup>** | 1 s.d. increase in advantage = 15.93% increase in probability of attracting decider |

In Galaxy group, a set’s success in attracting deciders was more likely when the total rate of acoustic signals given by its members was greater than the total rate of acoustic signals given by members of its opposing set (P1.1-2; β = 0.42, *p* = 4.5 x 10^-3^ ; table 2, figure 2d). As in Presidente group, deciders in Galaxy group were more likely to choose sets that had higher kinship advantages (P2.1; β = 0.85, *p* = 2.0 x 10^-5^; table 2, figure 2e) and higher frontness advantages (β = 0.96, *p* = 3.4 x 10^-5^; table 2; figure 2f).

While the tendency to follow the majority was stronger than the tendency to follow closer kin in Presidente group, members of Galaxy group had a stronger bias to follow closer kin than to follow the majority (P2.2; table 1). Despite this group-level difference, the strengths of the three biases we tested (toward majorities, kin, and frontal spatial positions) were broadly similar in both groups, in support of H2. For example, in Presidente group, increases of one standard deviation from the mean presence advantage, kinship advantage, and frontness advantage corresponded to increases in the predicted probability of attracting a decider by 24.48%, 12.20%, and 14.17%, respectively. In Galaxy group, increases of one standard deviation from the mean signal rate advantage, kinship advantage, and frontness advantage corresponded to increases in the predicted probability of attracting a decider by 11.26%, 16.97%, and 15.93%, respectively.

*H3: Instances of high directional disagreement allowed some individuals to exert social influence over collective movement decisions.* In both groups, the extent of interindividual variation in social influence was modest, with realized influence scores (calculated as an individual’s observed probability of winning an influence contest) ranging from 0.50 to 0.64 in Presidente group and from 0.41 to 0.61 in Galaxy group (table S5, figure 3a-b). As expected if resolutions to these conflicts were achieved via a shared decision-making process, neither group contained a single individual with a markedly higher probability of attracting others during instances of high directional conflict. The majority of Presidente group members (10/16; 62.50%) and Galaxy group members (7/11; 63.64%) won influence contests as frequently as expected under our null model of random decider choice. Nevertheless, six out of 16 Presidente group members (37.50%) and three out of 11 Galaxy group members (27.27%) were more likely to win influence contests than expected by chance (P3.1). Additionally, one of 11 Galaxy group members (9.09%) was less likely to win influence contests than expected by chance. Members of most age-sex classes present in Presidente group were represented among the individuals with higher-than-expected influence (except adult females). Conversely, only sub-adults had higher-than-expected influence in Galaxy group. The individual in Galaxy group with lower-than-expected influence was an adult female.

**Figure 3.**
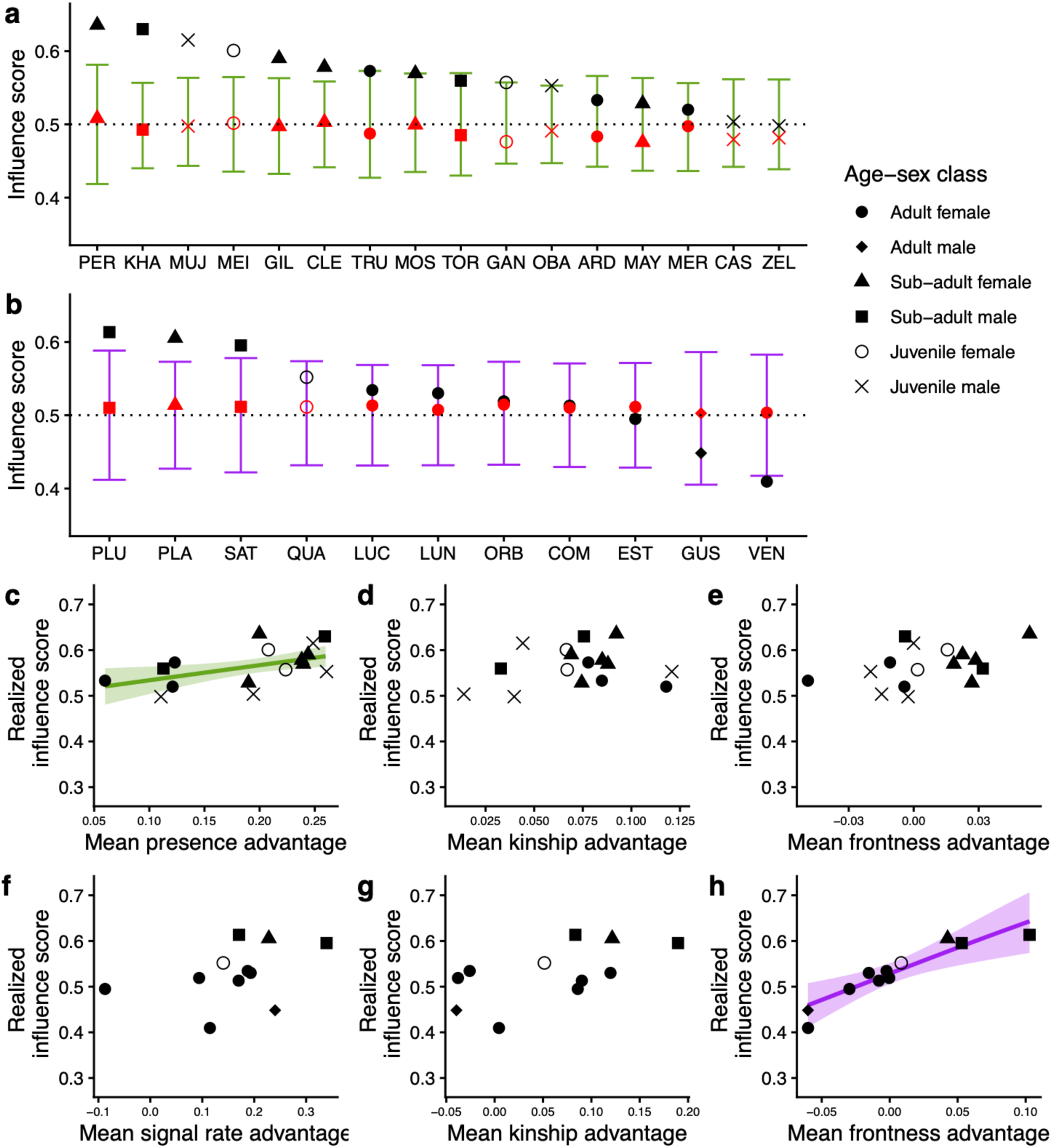
Social influence over travel direction varies across individuals. (a) and (b) show the distribution of influence scores in Presidente group and Galaxy group, respectively. Individuals are ordered by their realized influence scores (indicated by black points), from most to least influential. The lower and upper bounds of the bars for each individual indicate the 2.5^th^ percentile and 97.5^th^ percentile of the expected null distribution of realized influence scores if they had a random chance of winning influence contests. The red points indicate each individual’s baseline influence score (which statistically accounts for set-level advantages). Panels (c-e, Presidente group) and (f-h, Galaxy group) show the relationships between realized influence scores and individual mean presence (c) or signal rate (f) advantage, kinship advantage, and frontness advantage. Each point represents one individual and, where the association was statistically significant, the colored lines and shaded regions depict the model-predicted relationships and their 95% confidence intervals.

In both groups, interindividual variation in realized social influence was attributable to the biases of deciders investigated under H1-2. When statistically accounting for majority, kinship, and frontness advantages, interindividual variation in baseline social influence was virtually nonexistent (table S5, figure 3a-b). In Presidente group, baseline influence scores ranged from 0.48 to 0.51 and the random effect of individual participants was estimated to explain 0.10% of the total variance in influence contest outcomes. In Galaxy group, baseline influence scores ranged from 0.50 to 0.51 and the random effect of individual participants was estimated to explain 0.05% of the total variance in influence contest outcomes.

In Presidente group, realized social influence was correlated with the tendency to form majorities during directional conflicts: individuals with higher mean presence advantages had higher realized influence scores (P3.2; β = 1.34, *p* = 0.02; table S6, figure 3c). Across individuals, an increase of one standard deviation from the mean value of individual mean presence advantage predicted a 3.69% increase in realized influence score. However, social influence was not significantly predicted by mean kinship advantage (β = 0.55, *p* = 0.63; table S6, figure 3d) or mean frontness advantage (β = 1.61, *p* = 0.30; table S6, figure 3e).

In Galaxy group, spatial positioning was a stronger predictor of realized social influence: individuals with higher mean frontness advantages had higher realized influence during directional conflicts (P3.2; β = 4.61, *p* = 1.0 x 10^-3^; table S6, figure 3h). An increase of one standard deviation from the mean value of individual mean frontness advantage predicted a 10.37% increase in realized influence score. Social influence was not predicted by mean signal rate advantage (β = 0.10, *p* = 0.84; table S6, figure 3f) or mean kinship advantage (β = 0.41, *p* = 0.59; table S6, figure 3g).

## Discussion

When choosing between mutually exclusive travel directions, white-nosed coatis were biased by majority opinions, kin relationships, and spatial positioning and these biases produced moderate variation in realized social influence across individuals. This evidence suggests that complex social and demographic processes determine which individuals have outsized effects on collective decision outcomes in wild animal groups that make shared decisions.

We found some support for our first hypothesis. Coatis likely used majority-based heuristics to resolve these directional conflicts, but members of different groups were either more likely to choose directions that were indicated by a majority of total individuals (in Presidente group) or by a greater concentration of acoustic signals and individuals (in Galaxy group). This between-group difference may have several potential explanations. First, it may be that members of the different groups differed in their relative reliance on acoustic signals versus movement cues during directional conflicts, possibly because there were more juveniles (who emit contact calls at a lower rate than adults [38]) in Presidente group than in Galaxy group. Alternatively, members of both groups may have effectively integrated acoustic signals and movement cues to make accurate assessments about majority opinions, but the members of Galaxy group may have been less motivated to follow absolute majority opinions and maintain cohesion. This could be also related to between-group differences in age structure, or kinship (members of Galaxy group were generally less related than members of Presidente group). Furthermore, potential error exists in the automated detection of contact calls and in the automated designation of each call detection as coming from the animal wearing the collar versus a nearby conspecific (see Supplementary Methods). While we have no reason to expect that these sources of error should be biased to one group or another, it nevertheless remains possible that biased error in the contact call data set could contribute to between-group differences.

In support of our second hypothesis, the preferences of closer relatives were approximately as predictive of directional choice as were perceived majorities. This kin bias corroborates previous work in this system which demonstrated that coatis prioritize maintaining association with kin when groups split [34], and aligns with findings on collective movement dynamics in other female-philopatric species [51]. Notably, majority opinions were apparently less predictive of individual coati movement decisions during directional conflicts than has been reported in the literature for members of more socially cohesive groups, including guineafowl (*Acryllium vulturinum*), baboons (*Papio anubis*), and macaques (*Macaca tonkeana*) [9, 10, 13]. While this difference could reflect methodological differences across studies, it suggests that individual variation in responsiveness to majority opinion could be related to group- and population-level variation in social cohesion, either as a direct causal factor (e.g., groups become less cohesive as a by-product of low individual levels of responsiveness to majority opinions) or as a reactive behavioral response to the costs and benefits of cohesion (e.g., individuals become less attentive to majority opinions because the costs of cohesion increase).

In support of our third hypothesis, some individuals in both groups had higher social influence over group travel direction than was expected by chance, in that they were more likely to recruit groupmates toward the direction they preferred during directional conflicts. This result highlights that even when voting-like conflict resolution tactics are present, and obvious leaders are absent, individual variation in social influence over collective movement can still emerge when within-group dissimilarities in outcome preference are high. Studies of animal leadership often quantify the rate or probability of behavioral indicators of high social influence (e.g., being followed after initiating movement [52], high ordinal rank in travel progressions [53]) from observations sampled randomly across time. However, concentrating on instances when groupmates possess incompatible preferences could produce more biologically meaningful estimates of variation in control over where groups go.

Why did certain individuals have high social influence over travel direction? We found that interindividual variation in realized social influence was likely explained by variation in the same factors that biased the directional choices of deciders, given that interindividual variation in baseline social influence disappeared when statistically accounting for these decider biases. In Presidente group, an individual’s social influence score was positively predicted by its mean presence advantage. In other words, when disagreements over travel direction occurred, some coatis had greater tendencies to belong to majorities and also were more likely to recruit additional groupmates toward their preferred direction. This link between the tendency for majority formation and social influence mirrors the mechanism proposed for high social influence among male guineafowl [10, 17]. Galaxy group (in which deciders were generally less responsive to majority opinions than in Presidente group) exhibited a different pattern: an individual’s social influence score was positively predicted by its mean frontness advantage. In that group, three sub-adults had higher-than-expected social influence scores, forming an influential cluster who routinely signaled from more frontal sets and who likely were also all full siblings in a group with otherwise low relatedness. This pattern, in which a minority of group members with interdependent movement patterns form a unit with outsized influence over the group as a whole, resembles the higher-order structure of the influence network reported for a group of chacma baboons (*Papio ursinus*) [18]. We caution that the realized influence scores in a single group were likely not truly independent, as high influence individuals could mutually increase each other’s influence score by participating in sets together. This potential source of non-independence limits our ability for robust GLM-based estimation of the predictors of high influence in our current data set. However, support for H3 also comes from the negligible variation in baseline influence scores, which should not have been impacted by this within-group non-independence issue.

The structural advantages that some individuals receive from decider biases (e.g., toward majority opinions, kin, and frontward positions) likely emerge from complex, higher-order behavioral interactions, such that correlations between individual traits and social influence may be difficult to predict and highly dependent on group social structure and demography [17, 18, 54]. For example, Presidente group exhibited a ‘typical’ distribution of individuals in each age-sex class expected for white-nosed coati groups, while Galaxy group only had one surviving juvenile, and Trago group had no adult females. Which classes of individuals had high social influence also varied across these three groups: high influence individuals were spread across age-sex classes in Presidente group (figure 3a), concentrated solely among sub-adults in Galaxy group (figure 3b), and apparently absent in Trago group (table S5). While age is commonly cited as a source of individual variation in social influence [52, 55], our results suggest that such trait-based predictors of influence may be better understood when considered in light of group-level structure and demography.

Resolving conflicts of interest is a fundamental task for any group with heterogeneous membership. While some general strategies favoring conflict resolution are likely widespread across group-living taxa (e.g., conformity to majority opinions: [9, 10, 13]), they are combined with, and possibly counteracted by, individual-specific biases (e.g., conformity to opinions of close kin: this study). These biases that steer groups in one direction or the other can consistently favor the preferences of certain individuals, leading to inequality in social influence even when collective decisions are shared by most group members. Our results emphasize the particular importance of instances of high disagreement between groupmates to patterns of leadership and decision making. Moreover, the patterns shown across just two coati groups suggested that the distribution of social influence within groups is intimately tied to aspects of group demography and socio-spatial structure. In light of this, we anticipate that gaining a holistic understanding of how social influence over collective decisions is distributed within any system will require data on substantially more group replicates than is currently typical in the field, alongside advancements in theory.

## Author contributions

J.C.W.: Conceptualization, Data curation, Formal analysis, Investigation, Methodology, Visualization, Writing – original draft; E.M.G.: Conceptualization, Data curation, Investigation, Methodology, Project administration, Software, Writing – review & editing; J.O.: Data curation, Project administration, Writing – review & editing; M.J.Q.: Formal analyses, Writing – review & editing; M.C.C.: Conceptualization, Funding acquisition, Investigation, Project administration, Supervision, Writing – review & editing; B.T.H: Conceptualization, Data curation, Funding acquisition, Investigation, Methodology, Project administration, Supervision, Writing – review & editing; A.S.-P.: Conceptualization, Data curation, Funding acquisition, Investigation, Methodology, Project administration, Software, Supervision, Writing – review & editing

## Acknowledgements

We thank the Smithsonian Tropical Research Institute for permission to conduct research on BCI and in SNP, and MiAmbiente (permit number: SE/a-38-2020) and the Republic of Panama (permit number: PA-01-ARH-160-2022) for permission to export biological samples.

We also thank Lil Camacho for processing biological sample export permits, Patrick Paetzold for designing the housing of on-collar audio recorders, and Carolina Mitre Ramos, Brandol Ortega, and Lucía Torrez for assistance during capturing, collaring, and data collection. We are grateful to Julia Plecher, Leonie Treyer, Caroline Preuß, Roshni Keshwani, and Zeynep Durmuş for assistance with audio data labelling and verification and to Julian Schäfer-Zimmermann for his contributions and insight regarding automated vocalization detections. We are grateful to the members of the Communication and Collective Movement research group, Communication and Coordination Across Scales team, and the Department for the Ecology of Animal Societies at the Max Planck Institute of Animal Behavior for feedback on this project and to two anonymous reviewers for constructive comments on a previous version of this manuscript. A.S.P. and B.T.H. received funding for this research from Human Frontier Science Program Research Grant RGP0051/2019. A.S.P. received additional funding from the Gips-Schüle Stiftung and Deutsche Forschungsgemeinschaft under Germany’s Excellence Strategy – EXC 2117-422037984. Further funding was provided by M.C.C.’s Alexander von Humboldt Professorship (endowed by the Federal Ministry of Education and Research) and by the Max Planck Society.

