## Supplementary Material for "Kinship, acoustic signaling, and socio-spatial structure shape shared decisions and social influence in white-nosed coatis"

### Supplementary Methods

*Additional collaring details.* All collars were equipped with automated drop-off devices (Micro-TRD, Lotek, Newmarket, Canada) programmed to release after 18 days of tracking. Due to technical issues, all drop-off devices on collars deployed in Galaxy group and a minority of drop-off devices on collars deployed in Trago group failed to release on schedule. Therefore, we recaptured these individuals with failed drop-offs and manually removed the collars. All collars with failed drop-off devices were successfully removed within four weeks of the planned drop-off date. The drop-off mechanisms on two collars in Presidente group (affixed to sub-adults 'Moscoso' and 'Peron') released prematurely. One individual ('Moscoso') was recaptured, re-collared, and released within 5 hours of the premature drop-off and no data relevant for this study were lost (because the drop-off and recapture occurred after the daily GPS and audio sampling period ended, see below). The other individual ('Peron') was also recaptured and re-collared, but after a longer delay. As a result, there was a gap in GPS and audio data collection on this individual lasting 101 hours from start to finish.

*GPS and audio data processing.* Prior to analyses, we converted the GPS data to Universal Transverse Mercator (UTM) coordinates and performed minimal pre-processing to remove unrealistic locations and interpolate missing positions during short gaps (≤ 5 s). We also chose to exclude the first three days of data for one adult female in the Galaxy group ('Venus') because she exhibited unusually low levels of movement during this period, which immediately followed her initial capture and release.

We used the same audio processing protocol described in Grout *et al*. [1]. The same description can be found in that study's supplementary materials. The audio recorders generated a large data set of raw audio files (> 1,800 h). To identify and classify coati vocalizations from this data set, we trained a self-supervised transformer-based neural network model (*animal2vec*, [2]) on 115,760 manually-labelled vocalizations that were annotated from a subset of audio files in Adobe Audition v26.00.0.56 (Adobe Inc., San Jose, California). Label files consisted of contiguous sequences of audio for which the onsets and offsets of all calls within that segment were labelled. Each audio label encoded two classifications: (i) call type and (ii) focal status. The focal status of each call could either be classified as 'focal' (meaning the call was thought to be produced by the individual wearing the collar) or 'nonfocal' (meaning the call was thought to be produced by another individual within detection range of the recorder mounted on the focal individual's collar). Human labelers primarily distinguished between 'focal' and 'nonfocal' calls based on call amplitude and additional contextual information where available, but we note that these classifications are likely prone to greater error than classifications for call type. *animal2vec* was trained and tested using all 10-s audio clips for which at least one acoustic event was present, following Schäfer-Zimmerman *et al.* [2]. A subset of the manually-labelled calls (n = 28,970 calls) were excluded from the training set to serve as a validation set. With this validation set, we estimated a mean precision of 0.85, micro-averaged across call types .

After training *animal2vec* to detect coati calls and classify them by call type and focal status, we used it to generate call detections (onset time, offset time, call type, and focal status) for all audio files in our data set. *animal2vec* typically produces predictions on a fine-grained, frame-by-frame basis; here, we adopt the post-processing method described by the developers [2]. Event (i.e., call) boundaries are determined by applying a sliding average-pooling window to the likelihood output, which is subsequently binarized through a fixed threshold. The segment is then assigned a likelihood score as the mean of the frame-wise likelihoods within the predicted sequence. This workflow effectively converts frame-level data into instance-level metrics, resulting in a single prediction for each discrete event. The likelihood per event is bounded between 0 and 1, which indicates the model's confidence that the call belongs to the assigned class.

For each likelihood score, we assessed the recall (fraction of real calls of a given class that were correctly detected as that class) and precision (fraction of detected calls of a given class that actually belonged to that class). To assess the recall, we used the subset of data held out from training. However, because this hold-out set only included 10-second clips in which at least one acoustic event was detected, computing the precision based on this set would likely result in an underestimate, because a large proportion of the raw audio files consists of audio for which no calls are present but the machine learning model might still produce false detections. Therefore, to get an accurate assessment of precision, we generated predictions from *animal2vec* across 25 complete (3-hour) audio files that were previously not used in training, and that represented all individuals in Galaxy and Presidente groups (one file per individual), then manually verified the call type and focal status of the resulting 45,954 machine learning-generated detections. Using these verified files, we then computed the precision associated with each likelihood score based on how many detections were correct and how many were false positives. Finally, we combined these values with the associated recall score at each likelihood value to generate the precision-recall curve shown in figure S1.

For the purposes of this study, we were specifically interested in call types hypothesized to serve as 'contact calls': *chirp*, *chirp click*, *chirp grunt*, *click*, and *click grunt*. Therefore, we collapsed these acoustically distinguishable calls into one hypothesized functional category hereafter referred to as 'contact calls', and calculated the precision-recall curve for this 'contact call superclass'. Here, for each likelihood score, we defined the associated recall as the fraction of all (ground truth-labelled) focal calls of any contact call type that were detected by *animal2vec* as any contact call type (using the hold-out set). We defined the precision associated with a given likelihood score as the fraction of detections of any focal contact call type that truly belonged to any focal contact call category (using the verified files).

Based on these results, we excluded all contact call detections that were assigned a likelihood score of $\leq 0.40$. At this threshold, we estimated that contact calls were detected with a precision of 0.81 and a recall of 0.81 (figure S1).

*Determining subgroup compositions.* Because coati groups often split into foraging subgroups throughout the day, we had to determine the composition of each (sub)group at each time point. Following Della Libera et al. [3], (sub)group compositions were determined iteratively, starting by establishing 'connections' between every coati dyad that was separated by less than 15 m. Clusters of connected coatis were considered members of the same (sub)group, akin to the 'chain rule' sometimes used to observationally define group co-membership in the field. Applying a single distance threshold to define group co-membership to data collected with GPS collars can result in rapid fluctuations in group composition, because short changes in interindividual distance (e.g., from 14.5 m to 15.5 m) can break or make connections. Therefore, the sticky DBSCAN method applies a second, larger radius to define when connections are formally made and broken. A period of dyadic connection started at the first time point when two individuals were within a distance less than the larger radius (50 m), but was only counted if the pair eventually went within the smaller radius (15 m) before returning to a distance greater than the larger radius. A period of connection ends at the first time point when the distance between them was greater than the larger radius.

*Detecting influence contests.* We iterated through each individual in our data set to detect times at which they acted as a decider in an influence contest. For each coati $i$ at each time $t$, we first identified every groupmate $j$ who was 10 to 30 m away from $i$ and whose contact call rate was > 0.00 calls/s. We then defined each vector $\overrightarrow{ij}$ pointing from $i$ to one of these signaling groupmates and calculated the angle between every possible pair of vectors $\overrightarrow{ij}$. If the greatest of these angles was > 90°, then we considered that, from $i$'s perspective, the two groupmates forming the greatest angle (groupmate $j_A$ and groupmate $j_B$) were potentially indicating alternate future directions of travel. We then identified every other groupmate who was located, from $i$'s perspective, within 45° of $j_A$ or $j_B$ and < 30 m away. Together with groupmates $j_A$ and $j_B$, respectively, these two collections of conspecifics constituted set $A$ and set $B$. We chose 10 m as a lower proximity threshold for potential directional indicators because we assumed that some spatial separation was necessary to elicit following by a decider and we chose 30 m as an upper bound because we assumed that the ability to accurately detect and locate conspecifics would decline as a function of distance. We chose a 90° angle between potential travel directions as a threshold for substantial disagreement because this approximates the disagreement angle at which directional conflict strategies typically transition from preference averaging to choosing one preferred direction or the other [4, 5].

Within these times at which individual $i$ had two opposing potential travel directions, we aimed to isolate instances in which $i$ was initially stationary and then moved toward one of the two potential travel directions. Therefore, we excluded all times at which $i$'s speed was ≥ 0.05 m/s. Next, at each time $t$ when $i$'s speed was < 0.05 m/s and $i$ had two sets of individuals indicating alternate travel directions, we calculated the centroid of all individuals in each set, $A$ and $B$, and defined two new vectors: one pointing from $i$ to the centroid of set $A$ ($\overrightarrow{iA}$) and one pointing from $i$ to the centroid of set $B$ ($\overrightarrow{iB}$). We projected two artificial 'finish lines' that were both 10 m from *i*'s position; one perpendicularly crossing vector $\overrightarrow{iA}$ and one perpendicularly crossing vector $\overrightarrow{iB}$. Then, we determined whether *i* crossed either finish line in the next 200 s. If *i* never crossed either finish line by *t* + 200 s, then no influence contest was considered to have occurred at time *t*. We extracted all periods of consecutive time steps at which the time until *i* crossed a finish line was < 200 s. Whenever one of these periods began > 200 s after the end of the previous period, the first time in the period was recorded as the start time of an influence contest. The set associated with the first finish line crossed by *i* after this start time was considered to have 'won' the contest. Thus, each influence contest represents an event during which two sets of coatis were vocally signaling from different directions relative to another stationary coati, who then moved at least 10 m in one of those directions at > 0.2 m/s.

If detected influence contests contributed to collective movement decisions at a biologically meaningful scale, then we expected that, after contests occurred, entire groups (not just deciders) would be more likely to move in the directions indicated by winning sets than the directions indicated by losing sets. Therefore, at 20-s intervals over the hour following each influence contest, we calculated the centroid of the positions of all non-decider individuals who were present for the contest. We then defined a vector at each 20-s interval that pointed from their centroid at the start time of the influence contest to their current centroid. We calculated the angular difference between each of these future group heading vectors and $\overrightarrow{iA}$, and between each future group heading vector and $\overrightarrow{iB}$. We predicted that, on average, future group heading vectors would be more aligned (smaller angular difference) with the vector ($\overrightarrow{iA}$ or $\overrightarrow{iB}$) that connected the decider to the winner than the vector that connected the decider to the loser. This prediction held true: after influence contests occurred, group headings were, on average, more closely aligned with the vector pointing from the decider to the centroid of the winning set than with the vector pointing from the decider to the centroid of the losing set (figure S2).

*Calculating contact call rates.* Once contact call detections were time-matched to GPS locations, we were able to calculate rates of contact calling for each individual at all times when both GPS and audio data were collected. At time *t*, an individual *i*'s contact calling rate was calculated as the total number of contact calls that were detected on *i*'s audio recorder and assigned focal status over the period lasting from *t* – 99.5 s to *t* + 99.5 s, divided by 200. We calculated these rates over 200-s windows because 200 s was approximately the median length of time it took a coati to move 10 m (the distance over which we considered individual movement decisions, see *Defining influence contests* in Main Text) during the tracking windows.

*Presence, signaler, and signal advantage.* During directional conflicts, animals may rely on conspecific acoustic signals, cues about movement (e.g., visual, olfactory, or auditory indicators of neighbor locations and travel directions), or both to determine which direction most of their groupmates are going [6]. To account for these factors, we calculated three proxies of relative numerical support for both potential travel directions in every influence contest to assess the extent to which coatis follow majority opinions. First, we calculated the 'presence advantage' held by each set of influence contest participants as

$$\frac{(total\ individuals\ in\ focal\ set - total\ individuals\ in\ opposing\ set)}{(total\ individuals\ in\ focal\ set + total\ individuals\ in\ opposing\ set)}$$

This value ranged from -1 to 1, with negative numbers indicating that the set had fewer members than its opponent and positive numbers indicating that the set had more members than its opponent. In the example depicted in figure 1, set A had the greater presence advantage. Next, we calculated each set's 'signaler count advantage'. We used the same equation as for presence advantage, but substituted the total number of individuals in the set whose contact call rate at the time of the contest was > 0.00 calls/s in place of the total number of individuals contained in the set (in other words, we only counted individuals who were producing contact calls). In figure 1, sets A and B had equal signaler count advantages. Third, we determined each set's 'signal rate advantage'. This was again calculated in the same way as the other majority proxies, but was based on the relative difference between the sums of the call rates of all individuals in each set, rather than on the relative difference between the number of individuals present or vocalizing in each set. Set B in figure 1 had a greater signal rate advantage than set A.

*Kinship advantage.* To assess whether kinship predicted coati directional decisions, we used tissue samples collected during capture to determine relatedness estimates for every dyad via single nucleotide polymorphism genotyping. See Grout et al. [7] for full details on generating dyadic relatedness estimates. Within each set of participants in an influence contest, we determined which signaling member was most closely related to the decider and what their relatedness coefficient ($r$) to the decider was. We then calculated a set's 'kinship advantage' as

$$\frac{(maximum\ r\ in\ focal\ set - maximum\ r\ in\ opposing\ set)}{(maximum\ r\ in\ focal\ set + maximum\ r\ in\ opposing\ set)}$$

In figure 1, set A had a greater kinship advantage than set B because the decider's closest relative in set A was a closer relative than the decider's closest relative in set B. We chose to consider differences between the maximum relatedness of opposing sets to the decider (rather than mean relatedness of the decider to the members of each set) because we expected that this would be a less cognitively demanding decision rule for deciders to follow. Nevertheless, advantages calculated from differences in maximum relatedness and advantages calculated from differences in mean relatedness were highly correlated (Pearson's correlation coefficient = 0.90; $p < 2.2 \times 10^{-16}$), such that we do not expect that our results would differ meaningfully based on this decision.

*Frontness advantage.* We expected that individual directional decisions would be biased toward groupmates who were farther in front of the decider for several reasons. First we expected that coatis would be biased against tight turns because the turn angles animals exhibit while moving are not randomly distributed [8]. Second, we thought that groupmates who were in front of deciders might be more detectable that groupmates who were behind deciders (given coatis' rostrally-directed eyes, ears, and noses). Finally, because coatis in more frontal spatial positions have higher foraging success at fruit patches [9], deciders should be motivated to maintain travel pace with potential within-group competitors who are farther in front of them (even relative to other individuals who are also in front of them). Therefore, we calculated the 'frontness advantage' of each set of influence contest participants relative to both the previous heading of the decider's trajectory and to the decider's current position. Note that this metric is calculated from the perspective of the decider, and differs from group-level measures of frontness reported in previous studies on coati spatial positioning [9, 10]. We first defined the decider's previous heading as the vector pointing from its most recent location that was at least 10 m away from its current location to its current location (red dotted line in figure 1). We then took the centroid of the positions of a set's participants and projected it onto a coordinate plane defined with the decider's current position at the origin and its past heading as the y-axis. The set's frontness was defined as the centroid's position along the y-axis of this plane (positive values indicate the set was in front of the decider and negative values indicate the set was behind the decider). Unlike relatedness coefficients, counts of individuals, and call rates, frontness values included negative numbers. Therefore, we normalized all frontness values of sets *s* in group *g* on a scale from 0 to 1 with the equation

$$\frac{(frontness\ of\ set\ s - min.frontness\ value\ observed\ in\ group\ g)}{(max.frontness\ value\ observed\ in\ group\ g - min.frontness\ value\ observed\ in\ group\ g)}$$

before calculating frontness advantage as

$$\frac{(normalized\ frontness\ of\ focal\ set - normalized\ frontness\ of\ opposing\ set)}{(normalized\ frontness\ of\ focal\ set + normalized\ frontness\ of\ opposing\ set)}$$

In figure 1, set B had a greater frontness advantage than set A.

### Supplementary Results

*Summary statistics regarding detected influence contests.* In Presidente group, each set of participants in an influence contest included an average of 4.39 ± 2.48 total individuals (mean ± s.d.) and 3.65 ± 2.18 acoustic signalers. In Galaxy group, each set included a mean of 2.66 ± 1.59 total individuals and 2.39 ± 1.41 acoustic signalers. All individuals acted as deciders and participants in multiple influence contests. In Presidente group, individuals were deciders for

99.75 ± 21.25 influence contests, while this value was 66.55 ± 18.96 contests in Galaxy group. In the down-sampled subset of contests used to calculate participant-based metrics, individuals in Presidente group participated in 243.88 ± 45.43 contests and individuals in Galaxy group participated in 169.27 ± 32.30 contests.

Influence contests generally resulted in a consensus decision about travel direction, rather than a group fission. Five minutes after an influence contest occurred, the members of winning and losing sets were still together in the same subgroup 88.56% of the time in Presidente group and 86.15% of the time in Galaxy group. After influence contests occurred, group headings were, on average, more closely aligned with the vector pointing from the decider to the centroid of the winning set than with the vector pointing from the decider to the centroid of the losing set (figure S2). This difference persisted for up to an hour after an influence contest occurred, suggesting that these contest outcomes reflect biologically meaningful decisions about group travel direction.

*H1-2: Trago group results*. For Trago group data, the signaler count advantage model had the lowest AIC value and was a better fit to the data than the presence advantage model ($\Delta$AIC = 5.18) and the signal rate advantage model ($\Delta$AIC = 10.00). The results of the Trago presence advantage model, signaler advantage model, and signal advantage model are given in tables S7-9.

In Trago group, a given set was more likely to successfully attract a decider when it had relatively more vocal signalers than its opponent set ($\beta = 3.31$, $p = 5.6 \times 10^{-4}$; table S8) and when it was signaling from a relatively more frontward position (relative to the decider's past heading; $\beta = 1.50$, $p = 0.03$). A set's relative kinship advantage was not significantly correlated with its probability of attracting a decider ($\beta = -2.46$, $p = 0.08$).

*H3: Trago group results*. The realized influence scores of Trago group members ranged from 0.49 to 0.68, with no individuals having a realized influence score that was higher or lower than expected by chance (0/7 = 0.00%; table S5). Baseline influence scores showed no variation (range = [0.63, 0.63], table S5) and participant identity was estimated to explain 0.00% of the variance in influence contest outcome (but note that the small sample size for Trago group limits the interpretability of these estimates). Realized influence scores were not significantly predicted by mean signaler count advantage ($\beta = 1.19$, $p = 0.70$; table S10), kinship advantage ($\beta = 0.03$, $p = 0.99$), or frontness advantage ($\beta = 2.90$, $p = 0.25$).

### Supplementary Tables

*Table S1. Presidente group signaler count advantage model*. Results of a Bernoulli generalized linear mixed model predicting the probability that a randomly selected set of participants in an influence contest successfully attracted a decider in Presidente group (n = 1,596 decisions by 16 individuals). Statistically significant fixed effects are indicated in bold. A signed, multi-membership random effect matrix coding for the unique compositions of individuals present in the focal and nonfocal sets was included. To aid interpretation of fixed effects, we indicate the model-predicted percent change in the response variable with an increase of one standard deviation from the mean value of each fixed effect.

| Fixed effect | Estimate | Std. Error | *p*-value | Interpretation |
|---|---|---|---|---|
| Intercept | -0.03 | 0.05 | 0.56 | – |
| **Focal signaler count advantage** | **0.91** | **0.17** | **$1.2 \times 10^{-7}$** | 1 s.d. increase in advantage = 19.02% increase in probability of attracting decider |
| **Focal kinship advantage** | **0.76** | **0.18** | **$2.6 \times 10^{-5}$** | 1 s.d. increase in advantage = 12.67% increase in probability of attracting decider |
| **Focal frontness advantage** | **0.95** | **0.18** | **$4.8 \times 10^{-7}$** | 1 s.d. increase in advantage = 13.86% increase in probability of attracting decider |

*Table S2. Presidente group signal rate advantage model*. Results of a Bernoulli generalized linear mixed model predicting the probability that a randomly selected set of participants in an influence contest successfully attracted a decider in Presidente group (n = 1,596 decisions by 16 individuals). Statistically significant fixed effects are indicated in bold. A signed, multi-membership random effect matrix coding for the unique compositions of individuals present in the focal and nonfocal sets was included. To aid interpretation of fixed effects, we indicate the model-predicted percent change in the response variable with an increase of one standard deviation from the mean value of each fixed effect.

| Fixed effect | Estimate | Std. Error | *p*-value | Interpretation |
|---|---|---|---|---|
| Intercept | -0.03 | 0.05 | 0.53 | – |
| **Focal signal rate advantage** | **0.46** | **0.14** | **$1.2 \times 10^{-3}$** | 1 s.d. increase in advantage = 12.23% increase in probability of attracting decider |
| **Focal kinship advantage** | **0.96** | **0.19** | **$3.2 \times 10^{-7}$** | 1 s.d. increase in advantage = 14.46% increase in probability of attracting decider |
| **Focal frontness advantage** | **0.84** | **0.18** | **$2.4 \times 10^{-6}$** | 1 s.d. increase in advantage = 14.47% increase in probability of attracting decider |

*Table S3. Galaxy group presence advantage model*. Results of a Bernoulli generalized linear mixed model predicting the probability that a randomly selected set of participants in an influence contest successfully attracted a decider in Galaxy group (n = 732 decisions by 11 individuals). Statistically significant fixed effects are indicated in bold. A signed, multi-membership random effect matrix coding for the unique compositions of individuals present in the focal and nonfocal sets was included. To aid interpretation of fixed effects, we indicate the model-predicted percent change in the response variable with an increase of one standard deviation from the mean value of each fixed effect.

| Fixed effect | Estimate | Std. Error | *p*-value | Interpretation |
|---|---|---|---|---|
| Intercept | 0.04 | 0.08 | 0.60 | – |
| **Focal presence advantage** | **0.54** | **0.20** | **0.01** | 1 s.d. increase in advantage = 10.44% increase in probability of attracting decider |
| **Focal kinship advantage** | **0.82** | **0.20** | **$4.2 \times 10^{-5}$** | 1 s.d. increase in advantage = 16.44% increase in probability of attracting decider |
| **Focal frontness advantage** | **0.95** | **0.23** | **$3.6 \times 10^{-5}$** | 1 s.d. increase in advantage = 15.77% increase in probability of attracting decider |

*Table S4. Galaxy group signaler count advantage model*. Results of a Bernoulli generalized linear mixed model predicting the probability that a randomly selected set of participants in an influence contest successfully attracted a decider in Galaxy group (n = 732 decisions by 11 individuals). Statistically significant fixed effects are indicated in bold. A signed, multi-membership random effect matrix coding for the unique compositions of individuals present in the focal and nonfocal sets was included. To aid interpretation of fixed effects, we indicate the model-predicted percent change in the response variable with an increase of one standard deviation from the mean value of each fixed effect.

| Fixed effect | Estimate | Std. Error | *p*-value | Interpretation |
|---|---|---|---|---|
| Intercept | 0.05 | 0.08 | 0.56 | – |
| **Focal signaler count advantage** | **0.58** | **0.22** | **0.01** | 1 s.d. increase in advantage = 10.82% increase in probability of attracting decider |
| **Focal kinship advantage** | **0.80** | **0.20** | **$7.5 \times 10^{-5}$** | 1 s.d. increase in advantage = 16.01% increase in probability of attracting decider |
| **Focal frontness advantage** | **0.96** | **0.23** | **$2.7 \times 10^{-5}$** | 1 s.d. increase in advantage = 15.98% increase in probability of attracting decider |

*Table S5. Influence scores for all individuals*. Realized influence scores were calculated as the observed probability of winning an influence contest (i.e., attracting a conspecific during an instance of high directional conflict). Null distributions were generated for each individual under the assumption that deciders chose directions randomly with respect to which direction that individual indicated. Individuals whose realized influence scores were higher than expected by chance are indicated in bold and individuals whose realized influence scores were lower than expected by chance are italicized. Baseline influence scores were calculated as the model-predicted probability that an individual wins an influence contest, holding set-level advantages constant at zero.

| Group | Individual | Sex | Age class | Realized influence score | *Lower null bound* | *Upper null bound* | Baseline influence score |
|---|---|---|---|---|---|---|---|
| Presidente | Ardern | female | adult | 0.53 | 0.44 | 0.57 | 0.48 |
| | Castro | male | juvenile | 0.50 | 0.44 | 0.56 | 0.48 |
| | **Cleopatra** | **female** | **sub-adult** | **0.58** | **0.44** | **0.56** | **0.50** |
| | Gandhi | female | juvenile | 0.56 | 0.45 | 0.56 | 0.48 |
| | **Gillard** | **female** | **sub-adult** | **0.59** | **0.43** | **0.56** | **0.50** |
| | **Khan** | **male** | **sub-adult** | **0.63** | **0.44** | **0.56** | **0.49** |
| | May | female | sub-adult | 0.53 | 0.44 | 0.56 | 0.48 |
| | **Meir** | **female** | **juvenile** | **0.60** | **0.44** | **0.56** | **0.50** |
| | Merkel | female | adult | 0.52 | 0.44 | 0.56 | 0.50 |
| | Moscoso | female | sub-adult | 0.57 | 0.43 | 0.57 | 0.50 |
| | **Mujica** | **male** | **juvenile** | **0.62** | **0.44** | **0.56** | **0.50** |
| | Obama | male | juvenile | 0.55 | 0.45 | 0.55 | 0.49 |
| | **Peron** | **female** | **sub-adult** | **0.64** | **0.42** | **0.58** | **0.51** |
| | Torrijos | male | sub-adult | 0.56 | 0.43 | 0.57 | 0.59 |
| | Truss | female | adult | 0.57 | 0.43 | 0.57 | 0.49 |
| | Zelenskyy | male | juvenile | 0.50 | 0.44 | 0.56 | 0.48 |
| Galaxy | Cometa | female | adult | 0.51 | 0.43 | 0.57 | 0.51 |
| | Estrella | female | adult | 0.49 | 0.43 | 0.57 | 0.51 |
| | Gus | male | adult | 0.45 | 0.41 | 0.59 | 0.50 |

| | | | | | | | |
|---|---|---|---|---|---|---|---|
| | Lucero | female | adult | 0.53 | 0.43 | 0.57 | 0.51 |
| | Luna | female | adult | 0.53 | 0.43 | 0.57 | 0.51 |
| | Orbita | female | adult | 0.52 | 0.43 | 0.57 | 0.51 |
| | **Planeta** | **female** | **sub-adult** | **0.61** | **0.43** | **0.57** | **0.51** |
| | **Pluto** | **male** | **sub-adult** | **0.61** | **0.41** | **0.59** | **0.51** |
| | Quasar | female | juvenile | 0.55 | 0.43 | 0.57 | 0.51 |
| | **Saturno** | **male** | **sub-adult** | **0.60** | **0.42** | **0.58** | **0.51** |
| | *Venus* | *female* | *adult* | *0.41* | *0.42* | *0.58* | *0.50* |
| Trago | Amarula | female | juvenile | 0.59 | 0.34 | 0.66 | 0.63 |
| | Bailey | male | sub-adult | 0.49 | 0.35 | 0.65 | 0.63 |
| | Cerveza | female | juvenile | 0.65 | 0.32 | 0.68 | 0.63 |
| | Limoncello | male | juvenile | 0.56 | 0.33 | 0.67 | 0.63 |
| | Sake | female | juvenile | 0.61 | 0.32 | 0.68 | 0.63 |
| | Tiger | female | juvenile | 0.58 | 0.33 | 0.67 | 0.63 |
| | Whisky | female | juvenile | 0.68 | 0.32 | 0.68 | 0.63 |

*Table S6. Results of binomial GLMs predicting the proportion of influence contests that an individual won in Presidente group (n = 16 individuals) and Galaxy group (n = 11 individuals)*. Statistically significant fixed effects are indicated in bold.

| Group | Fixed effect | Estimate | Std. Error | *p*-value | Interpretation |
|---|---|---|---|---|---|
| Presidente | (intercept) | -0.05 | 0.13 | 0.72 | – |
| | **Mean presence advantage** | **1.34** | **0.55** | **0.02** | 1 s.d. increase in mean advantage = 3.69% increase in social influence score |
| | Mean kinship advantage | 0.55 | 1.14 | 0.63 | – |
| | Mean frontness advantage | 1.61 | 1.55 | 0.30 | – |
| Galaxy | (intercept) | 0.07 | 0.10 | 0.44 | – |
| | Mean signal rate advantage | 0.10 | 0.49 | 0.84 | – |
| | Mean kinship advantage | 0.41 | 0.76 | 0.59 | – |
| | **Mean frontness advantage** | **4.61** | **1.40** | **$1.0 \times 10^{-3}$** | 1 s.d. increase in mean advantage = 10.37% increase in social influence score |

*Table S7. Trago group presence advantage model*. Results of a Bernoulli generalized linear mixed model predicting the probability that a randomly selected set of participants in an influence contest successfully attracted a decider in Trago group (n = 73 decisions by 7 individuals). Statistically significant fixed effects are indicated in bold. A signed, multi-membership random effect matrix coding for the unique compositions of individuals present in the focal and nonfocal sets was included. To aid interpretation of fixed effects, we indicate the model-predicted percent change in the response variable with an increase of one standard deviation from the mean value of each fixed effect.

| Fixed effect | Estimate | Std. Error | *p*-value | Interpretation |
|---|---|---|---|---|
| Intercept | 0.50 | 0.28 | 0.07 | – |
| **Focal presence advantage** | **2.40** | **0.82** | **$3.6 \times 10^{-3}$** | 1 s.d. increase in advantage = 35.16% increase in probability of attracting decider |
| Focal kinship advantage | -1.71 | 1.32 | 0.20 | – |
| **Focal frontness advantage** | **1.23** | **0.62** | **0.05** | 1 s.d. increase in advantage = 21.09% increase in probability of attracting decider |

*Table S8. Trago group signaler count advantage model*. Results of a Bernoulli generalized linear mixed model predicting the probability that a randomly selected set of participants in an influence contest successfully attracted a decider in Trago group (n = 73 decisions by 7 individuals). Statistically significant fixed effects are indicated in bold. A signed, multi-membership random effect matrix coding for the unique compositions of individuals present in the focal and nonfocal sets was included. To aid interpretation of fixed effects, we indicate the model-predicted percent change in the response variable with an increase of one standard deviation from the mean value of each fixed effect.

| Fixed effect | Estimate | Std. Error | *p*-value | Interpretation |
|---|---|---|---|---|
| Intercept | 0.55 | 0.29 | 0.06 | – |
| **Focal signaler count advantage** | **3.31** | **0.96** | **$5.6 \times 10^{-4}$** | 1 s.d. increase in advantage = 41.18% increase in probability of attracting decider |
| Focal kinship advantage | -2.46 | 1.42 | 0.08 | – |
| **Focal frontness advantage** | **1.50** | **0.68** | **0.03** | 1 s.d. increase in advantage = 24.97% increase in probability of attracting decider |

*Table S9. Trago group signal rate advantage model*. Results of a Bernoulli generalized linear mixed model predicting the probability that a randomly selected set of participants in an influence contest successfully attracted a decider in Trago group (n = 73 decisions by 7 individuals). Statistically significant fixed effects are indicated in bold. A signed, multi-membership random effect matrix coding for the unique compositions of individuals present in the focal and nonfocal sets was included. To aid interpretation of fixed effects, we indicate the model-predicted percent change in the response variable with an increase of one standard deviation from the mean value of each fixed effect.

| Fixed effect | Estimate | Std. Error | *p*-value | Interpretation |
|---|---|---|---|---|
| Intercept | 0.35 | 0.26 | 0.18 | – |
| **Focal signal rate advantage** | **1.03** | **0.48** | **0.03** | 1 s.d. increase in advantage = 22.41% increase in probability of attracting decider |
| Focal kinship advantage | 0.11 | 1.03 | 0.91 | – |
| Focal frontness advantage | 1.08 | 0.59 | 0.07 | – |

*Table S10. Trago group individual influence model*. Results of a binomial generalized linear model predicting the proportion of influence contests that individuals in Trago group won (n = 7 individuals). No fixed effects were statistically significant.

| Fixed effect | Estimate | Std. Error | *p*-value | Interpretation |
|---|---|---|---|---|
| Intercept | 0.16 | 0.45 | 0.72 | – |
| Mean signaler count advantage | 1.19 | 3.07 | 0.70 | – |
| Mean kinship advantage | 0.03 | 2.80 | 0.99 | – |
| Mean frontness advantage | 2.90 | 2.50 | 0.25 | – |

**Supplementary Figures**

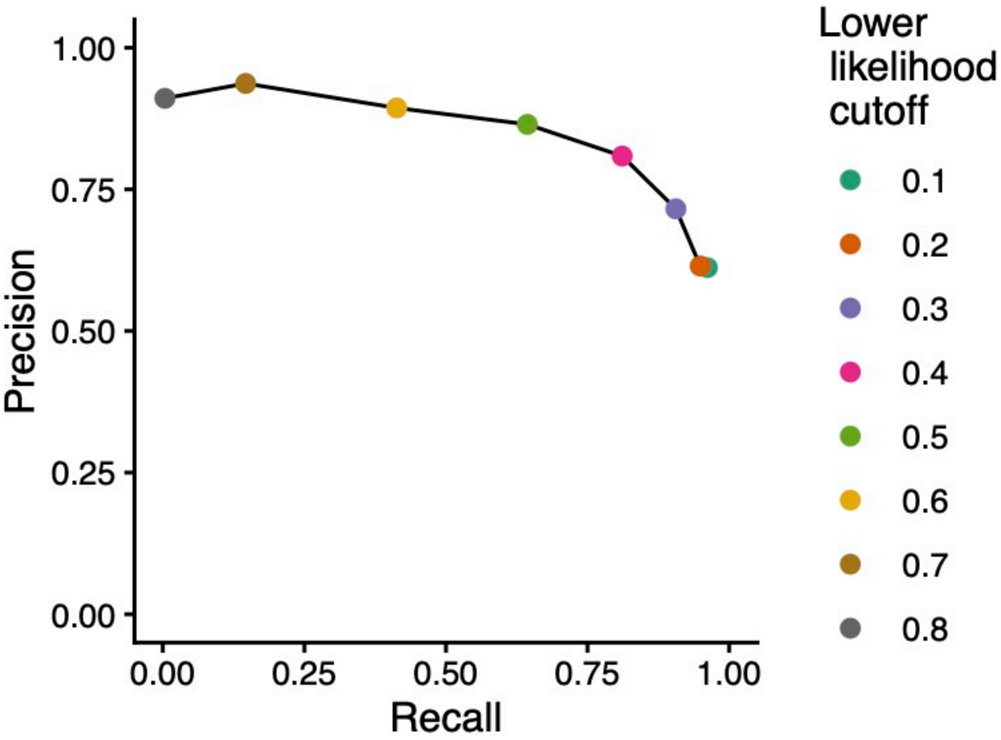

*Figure S1. Estimated precision and recall of contact call detections as a function of likelihood score*. A likelihood score was calculated for each machine learning-detected coati call. Each point represents the estimated precision and recall of contact call detections assigned a likelihood score greater than a given cutoff. Based on these estimates, we only analyzed detected contact calls that were assigned a likelihood score of > 0.40 (pink point) to optimally balance precision and recall. This led to a recall of 0.81 (81% of true contact calls detected) and a precision of 0.81 (81% of detected contact calls were correct).

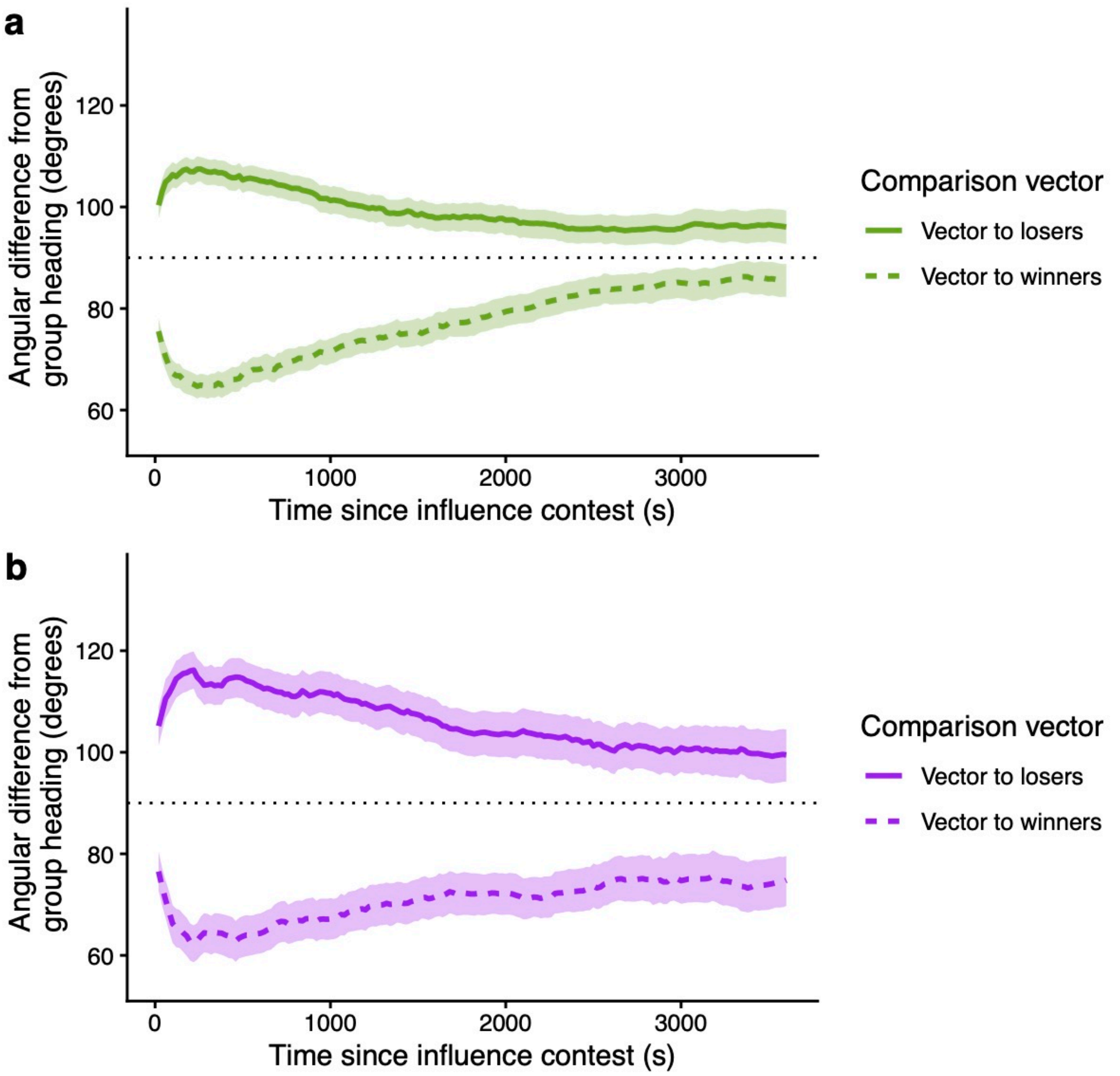

*Figure S2. Time-lagged correlations between directional options in influence contests and subsequent group-level movement*. Across all influence contests in Presidente (a) and Galaxy (b) groups, we calculated the average angular difference between the heading of the centroid of all non-decider individuals present at the contest and either the vector pointing from the contest decider to the winning set (dashed line) or the vector pointing from the contest decider to the losing set (solid line) over the hour following the contest. Shaded regions represent 95% confidence intervals of the mean based on nonparametric bootstrapping of the data sets. Larger angular differences indicate disagreement (the group did not ultimately move in that direction) and smaller angular differences alignment (the group ultimately moved in that direction). Under the null expectation, the confidence intervals would overlap 90° (indicated by the dotted black line). These patterns suggest that influence contests represent biologically meaningful decision points about collective travel direction.
